# Nuclease escape elements protect messenger RNA against cleavage by multiple viral endonucleases

**DOI:** 10.1101/155457

**Authors:** Mandy Muller, Britt A. Glaunsinger

**Author notes:** Present address: Department of Microbiology, University of Massachusetts at Amherst, Amherst, MA 01003 USA.

## Abstract

During lytic Kaposi’s sarcoma-associated herpesvirus (KSHV) infection, the viral endonuclease SOX promotes widespread degradation of cytoplasmic messenger RNA (mRNA). However, select mRNAs, including the transcript encoding interleukin-6 (IL-6), escape SOX-induced cleavage. IL-6 escape is mediated through a 3’ UTR RNA regulatory element that overrides the SOX targeting mechanism. Here, we reveal that this protective RNA element functions to broadly restrict cleavage by a range of homologous and non-homologous viral endonucleases. However, it does not impede cleavage by cellular endonucleases. The IL-6 protective sequence may be representative of a larger class of nuclease escape elements, as we identified a similar protective element in the GADD45B mRNA. The IL-6 and GADD45B-derived elements display similarities in their sequence, putative structure, and several associated RNA binding proteins. However, the overall composition of their ribonucleoprotein complexes appears distinct, leading to differences in the breadth of nucleases restricted. These findings highlight how RNA elements can selectively control transcript abundance in the background of widespread virus-induced mRNA degradation.

**AUTHOR SUMMARY:** The ability of viruses to control the host gene expression environment is crucial to promote viral infection. Many viruses express factors that reduce host gene expression through widespread mRNA decay. However, some mRNAs escape this fate, like the transcript encoding the immunoregulatory cytokine IL-6 during KSHV infection. IL-6 escape relies on an RNA regulatory element located in its 3’UTR and involves the recruitment of a protective protein complex. Here, we show that this escape extends beyond KSHV to a variety of related and unrelated viral endonucleases. However, the IL-6 element does not protect against cellular endonucleases, revealing for the first time a virus-specific nuclease escape element. We identified a related escape element in the GADD45B mRNA, which displays several similarities with the IL-6 element. However, these elements assemble a largely distinct complex of proteins, leading to differences in the breadth of their protective capacity. Collectively, these findings reveal how a putative new class of RNA elements function to control RNA fate in the background of widespread mRNA degradation by viral endonucleases.

## INTRODUCTION

A number of viruses restrict host gene expression to reduce competition for resources and dampen immune responses. This ‘host shutoff’ phenotype can be triggered through a range of mechanisms that operate at nearly every stage of the gene expression cascade. Viruses whose host shutoff strategies involve the induction of widespread mRNA decay include the alpha and gammaherpesviruses, vaccinia virus (VACV), influenza A virus (IAV), and SARS coronavirus (SCoV) [1-5] In each of the above cases, mRNA degradation is induced *via* one or more internal endonucleolytic cleavages in the target mRNA or, in the case of VACV, direct removal of the mRNA 5’ cap [5-9]. This is invariably followed by exonucleolytic degradation of the cleaved fragment(s) by components of the mammalian RNA decay machinery such as Xrn1 [1,10,11].

The viral strategy contrasts with basal mRNA degradation in eukaryotes, which is a tightly regulated process that initiates with gradual shortening of the poly(A) tail, followed by removal of the 5’ cap prior to exonucleolytic degradation of the transcript body [12]. Although eukaryotes encode endonucleases, they are generally restricted to a highly specific set of targets. For example, mRNAs containing premature stop codons are cleaved by the Smg6 endonuclease during the translation-linked quality control process of nonsense mediated decay (NMD) [13]. No-go decay is another form of quality control activated in cases of ribosome stalling, although the specific endonuclease that cleaves the mRNA remains unknown [14,15]. The use of endonucleases during quality control enables more rapid removal of aberrant mRNAs from the translation pool, as their inactivation is not reliant on the prior rate limiting steps of deadenylation and decapping. In this regard, virus-induced mRNA decay resembles the cellular quality control mechanisms, but with significantly expanded scope.

One of the well-studied viral endonucleases is the SOX protein encoded by ORF37 of Kaposi’s sarcoma-associated herpesvirus (KSHV). During lytic KSHV replication, SOX is expressed with delayed early kinetics and its nuclease activity significantly reduces cytoplasmic mRNA levels [16]. SOX is conserved throughout the herpesvirus family, but only gammaherpesviral SOX homologs display ribonuclease activity in cells [16-18]. Perhaps surprisingly, its activity is not restricted to host mRNAs, and studies with the SOX homolog from murine gammaherpesvirus 68 (MHV68) indicate that SOX activity helps fine tune viral mRNA levels in a manner important for the *in vivo* viral lifecycle [19,20]. Although most mRNAs are subject to cleavage by SOX, a recent degradome-based sequencing analysis together with studies on individual endogenous and reporter mRNAs revealed that SOX cleavage sites are defined by a degenerate RNA motif [10,21]. The SOX-targeting motif can be located anywhere within an mRNA and may be present multiple times [10,21].

The observation that a targeting motif is present on SOX cleaved mRNAs suggests that transcripts lacking this element should escape cleavage. Indeed, RNAseq analyses indicate that approximately one-third of mRNAs are not depleted by SOX [22,23]. Studying these ‘escapees’ in aggregate is complicated, however, by the fact that multiple mechanisms can promote apparent escape. These include lack of a targeting motif, indirect transcriptional effects, and active evasion of cleavage [22,24-28]. This latter phenotype, termed dominant escape, is particularly notable as it involves a specific RNA element whose presence in the 3’ UTR of an mRNA protects against SOX cleavage, regardless of whether the RNA contains a targeting motif. The one known example of dominant escape derives from the host interleukin-6 (IL-6) transcript [26,27,29].

IL-6 expression is required for survival of B cells infected with KSHV, and the virus engages a number of strategies to drive production of this cytokine [30-38]. The IL-6 mRNA is directly refractory to SOX cleavage and thus remains robustly induced during host shutoff due to the presence of a specific ‘SOX resistance element’ (SRE) [26,29]. Even in the absence of infection, reporter mRNAs bearing the IL-6 SRE remain stable in SOX-expressing cells, an observation that has helped delineate features of this novel RNA element required for the protective phenotype [26,27]. The IL-6 SRE was fine mapped to a 200 nt sequence within the 3’ UTR, which was subsequently shown to assemble an ‘escape complex’ of at least 8 cellular RNA binding proteins involved in protection [26,27]. How this complex functions to restrict SOX recognition remains largely unknown, although nucleolin (NCL) plays an essential role. NCL is partially relocalized from the nucleolus to the cytoplasm during lytic KSHV infection, where it binds the SRE using its RNA recognition motif and engages in protein-protein interactions related to escape with its carboxyl-terminal RGG domain [27].

The IL-6 SRE is the first described ribonuclease escape element and much remains to be learned about its function, as well as whether other related elements exist that protect their associated mRNA. Here, we reveal that the IL-6 SRE is broadly protective against a diverse group of viral endonucleases, suggesting an underlying commonality in the mechanism by which these host shutoff factors recognize their mRNA targets. It is not indiscriminately protective, however, as host quality control endonucleases are not blocked by the IL-6 SRE. We then identify a second, novel SRE within the GADD45B mRNA, which displays some physical and functional similarities to the IL-6 SRE. Collectively, these findings suggest that a diversity of nuclease escape elements exist, and that their characterization may lead to new insights into the control of mRNA fate in both infected and uninfected cells.

## RESULTS

### The IL-6 derived resistance element broadly protects against cleavage by viral but not cellular endonucleases

The 3’ UTR of IL-6 contains a transferrable 200 nt SRE that protects its associated mRNA from SOX-induced degradation [26,27]. The SRE also protects mRNA from cleavage by the unrelated vhs endonuclease encoded by herpes simplex virus type 1 (HSV-1), hinting that this RNA element may restrict endonuclease targeting in a broader capacity [27]. To test this hypothesis, we assessed whether the presence of the SRE impacted the ability of a panel of homologous and heterologous mRNA-specific viral endonucleases to degrade a target mRNA. In addition to HSV-1 vhs, these included homologs of KSHV SOX from the related gammaherpesviruses Epstein-Barr virus (EBV; BGLF5) and murine gammaherpesvirus 68 (MHV68; muSOX), as well as the heterologous host shutoff endonuclease from influenza A virus (IAV; PA-X) [3,17,39]. As anticipated, the control GFP mRNA was readily degraded in 293T cells co-transfected with plasmids expressing each of the viral endonucleases as measured by RT-qPCR (**Fig. 1A**). However, addition of the IL-6 derived SRE to the 3’ UTR of GFP (GFP-IL-6-SRE) prevented each of the viral endonucleases from degrading this mRNA (**Fig. 1B**). Fusion of a size matched segment of the IL-6 3’ UTR lacking the SRE to GFP (GFP-IL-6 ΔSRE) reinstated cleavage of the GFP reporter by the viral endonucleases (**Fig. 1B**). These data confirm that the protection conferred by the SRE is not specific to SOX-induced cleavage, but functions more broadly to restrict mRNA cleavage by a diverse set of mammalian virus host shutoff endonucleases.

**Figure 1:**
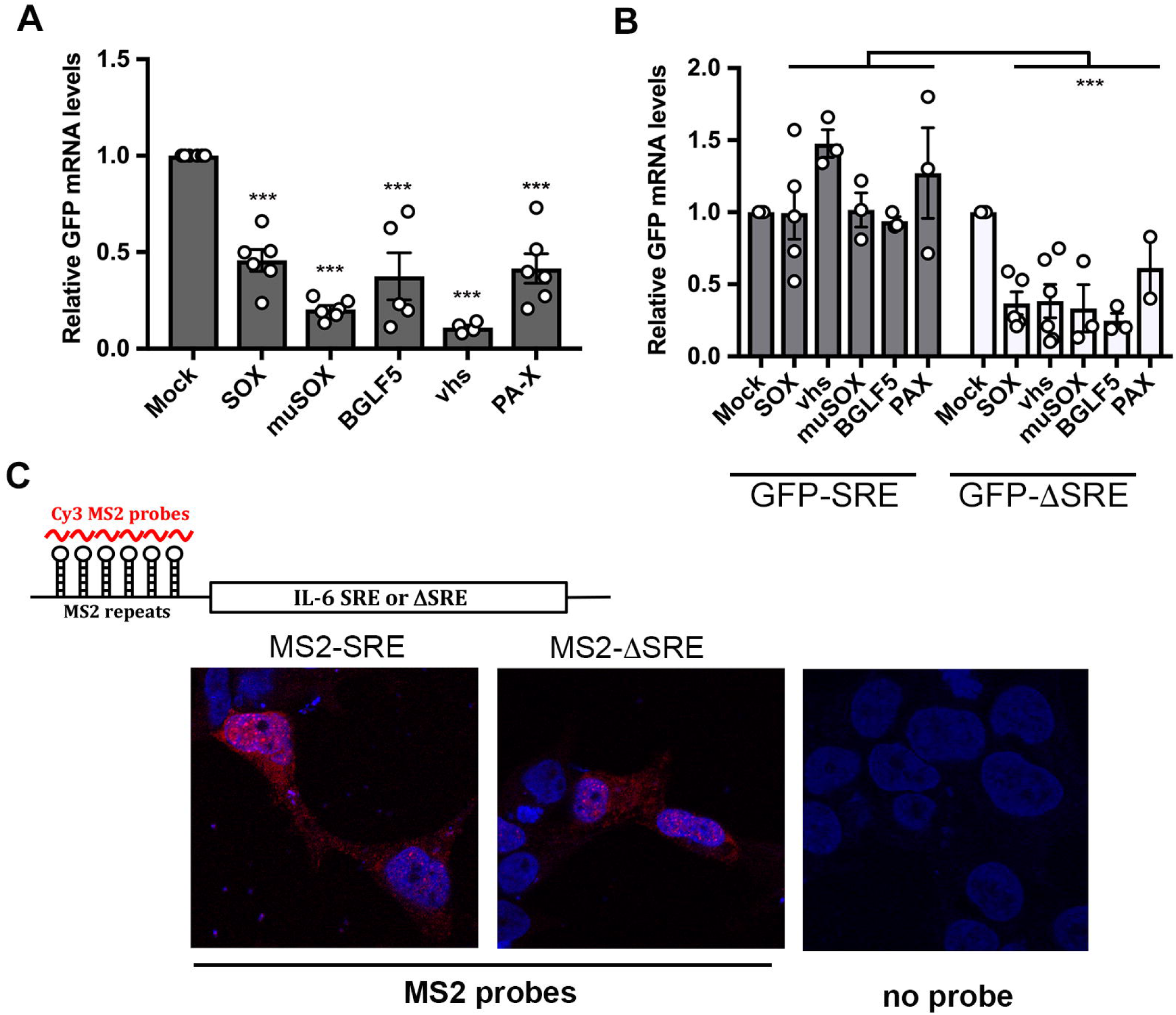
The IL-6-SRE broadly protects against multiple viral endonucleases. **(A)** 293T cells were transfected with empty vector (mock) or a plasmid expressing the indicated endonucleases along with a GFP reporter. After 24 h, total RNA was harvested and subjected RT-qPCR to measure GFP mRNA levels. Graphs here and afterwards display individual replicates as dots, together with the mean values (±SEM). Statistical significance was determined by the Student t test (* p<0.1; ** p<0.05; *** p<0.01). (**B**) 293T cells were co-transfected with the indicated endonuclease-expressing plasmid expressing together with a GFP-SRE or GFP-ΔSRE reporter. After 24 h, total RNA was harvested and subjected RT-qPCR to measure GFP mRNA levels. **(C)** Top: diagram showing the structure of the reporter mRNA containing MS2 repeats upstream of the SRE or ΔSRE fragment of the IL-6 3’UTR. Red lines denote the region detected by the MS2 probe. Bottom: 293T cells were transfected with the indicated MS2 reporters. 24h later, cells were fixed and processed for RNA FISH staining. Signals from the MS2 probes (red) and DAPI stained nuclei (blue) were detected by confocal microscopy.

Cleavage sites for IAV PA-X have yet to be determined, but the other endonucleases do not appear to target mRNA in the same location or using the same sequence features, including the SOX homologs [1,9,21,40,41]. Thus, the IL-6 derived SRE is unlikely to function through steric occlusion of a common cleavage site. We instead considered the possibility that the SRE functions like an RNA ‘zip code’, directing its associated transcript to a location in the cell inaccessible to endonucleases. To test this hypothesis, we used RNA fluorescence *in situ* hybridization (FISH) to monitor how the presence of the IL-6 SRE impacted the localization of its associated mRNA. Six phage MS2-derived stem-loops were introduced upstream of the SRE or ΔSRE segment of the IL-6 3’ UTR in the pcDNA3 luciferase reporter, enabling visualization of the RNA in transfected 293T cells using a Cy3-labeled RNA probe directed against the MS2 sequences [42]. We verified that the presence of the stem-loops did not prevent the escape of the SRE-containing reporter or degradation of the ΔSRE reporter in SOX-expressing cells (**Fig. S1**). There was no distinguishable difference in the localization of the SRE and ΔSRE containing mRNAs, both of which were present relatively diffusely throughout the cell (**Fig. 1C**). The FISH signal was specific to transfected cells, as we observed no fluorescence in neighboring untransfected cells nor autofluorescence from transfected cells lacking the Cy3 probes (**Fig. 1C**). Although these data to not exclude the possibility that the SRE-containing transcript became sequestered into micro-aggregates or other structures not visible at this level of resolution, they do not support relocalization as the driver of escape from viral endonuclease cleavage.

We next considered whether the inability to cleave an IL-6 SRE-containing transcript was specific to viral endonucleases or similarly extended to host endonucleases. Although basal cellular mRNA decay is carried out by exonucleases, host quality control pathways involve endonucleases to promote rapid clearance of aberrant mRNA [43-45]. We applied two strategies to monitor the activity of the SRE against host endonucleases. The first was to use a nonsense mediated decay (NMD) target containing a premature termination codon (PTC) 100 amino acids into the body of an RFP reporter mRNA (dsRed2-PTC) [10]. The NMD pathway detects PTCs during translation and directs cleavage of the mRNA by the Smg6 endonuclease [46]. Depletion of Smg6 from 293T cells using siRNAs restored dsRed2-PTC mRNA levels to those of the control dsRed2 transcript, confirming that this was an NMD substrate degraded by Smg6 (**Fig. 2A-B**). We then fused the IL-6 derived SRE to the 3’ UTR of dsRed2-PTC (dsRed2-PTC-SRE) or, as a control, the size matched region from the IL-6 3’ UTR lacking the SRE (dsRed2-PTC-ΔSRE) and measured the levels of each mRNA by RT-qPCR in 293T cells (**Fig. 2C**). In contrast to the viral endonucleases, the SRE did not impair Smg6-mediated cleavage of its target mRNA, as all three PTC-containing transcripts were similarly degraded (**Fig. 2A & 2C**).

**Figure 2:**
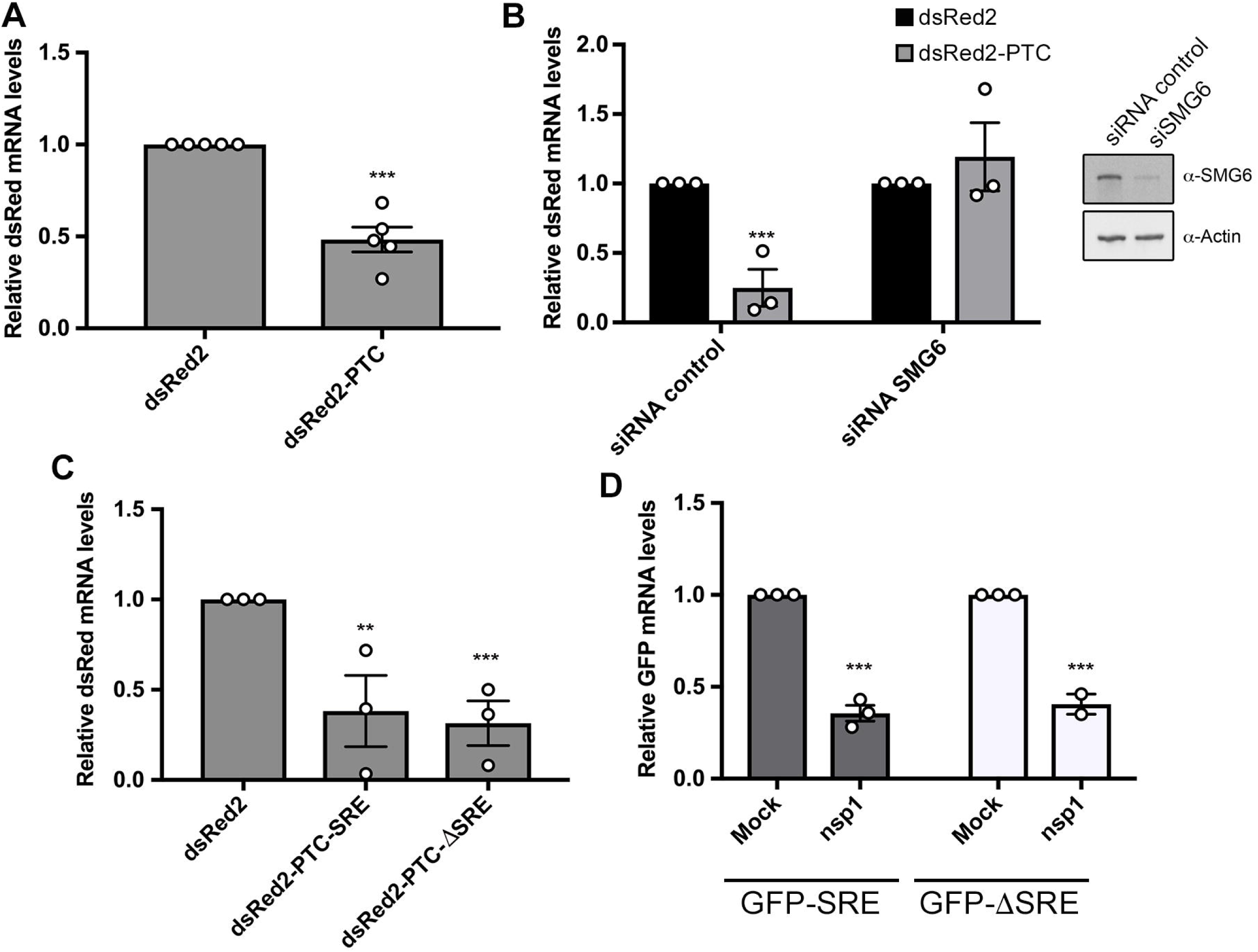
The IL-6-SRE does not protect against cellular endonucleases. **(A)** 293T cells were transfected with the indicated dsRed2 reporters. After 24 h, total RNA was harvested and subjected RT-qPCR to measure dsRed mRNA levels. **(B)** 293T cells were treated with siRNAs targeting Smg6 or control non-targeting siRNAs. 48h later, cells were transfected with the indicated dsRed reporters. After 24h, total RNA was harvested and subjected RT-qPCR to measure dsRed mRNA levels (*left*). The efficiency of SMG6 knockdown was measured by western blotting, with actin serving as a loading control (*right*). **(C)** 293T cells were transfected with the indicated dsRed2 reporters. After 24 h, total RNA was harvested and subjected to RT-qPCR to measure dsRed mRNA levels. **(D)** 293T cells were transfected with an empty vector (mock) or a plasmid expressing SCoV nsp1 along with the indicated GFP reporters. After 24 h, total RNA was harvested and subjected RT-qPCR to measure GFP mRNA levels.

The second strategy to monitor host endonuclease activity was to express the nsp1 protein from SARS coronavirus. Nsp1 is not itself a nuclease, but it binds the 40S ribosome and causes it to stall on the mRNA, thus activating cleavage of the mRNA by an as yet unknown cellular endonuclease via a mechanism reminiscent of no-go decay [47,48]. Although the endonuclease involved in this pathway has not been established, it is known not to be Smg6 and thus this enabled evaluation of a distinct host endonuclease [48]. Nsp1 was co-transfected with the GFP-SRE or GFP-ΔSRE reporter into 293T cells, and depletion of the GFP transcript was measured by RT-qPCR. Similar to the NMD substrate, the SRE did not prevent degradation of the GFP mRNA in nsp1-expressing cells (**Fig. 2D**). Collectively, these results suggest that the IL-6 derived SRE confers broad protection against viral but not cellular endonucleases. Given that the host endonucleases require ongoing translation for target recognition, these data also confirm that the SRE does not pull its associated mRNA out of the translation pool.

### GADD45B mRNA is also resistant to SOX-induced degradation

To determine whether this property of broad inhibition of viral endonucleases was restricted to the IL-6 SRE, we sought to identify other SRE-bearing transcript(s). Based on our previous finding that the IL-6 SRE required binding by a complex of cellular proteins including NCL, we mined a published RNAseq dataset for transcripts that were not depleted in SOX expressing 293T cells and were known to be bound by NCL [22,49-51]. A transcript that fit these criteria was growth arrest and DNA damage-inducible 45 beta (GADD45B). Notably, the GADD45B mRNA was also previously shown to escape degradation in HSV-1 vhs-expressing cells, further suggesting it might contain an SRE [52-54]. We first examined whether the GADD45B mRNA was resistant to host shutoff upon lytic reactivation of a KSHV-positive B cell line (TREX-BCBL1) and a renal carcinoma cell line stably expressing the KSHV BAC16 (iSLK.219). Both TREX-BCBL1 and iSLK.219 cells harbor a doxycycline (dox)-inducible version of the major viral lytic transactivator RTA that promotes entry into the lytic cycle upon dox treatment [55,56]. Unlike the GAPDH mRNA, which is degraded by SOX upon lytic reactivation, the GADD45B mRNA levels remained unchanged both in reactivated TREX-BCBL1 and iSLK.219 cells as measured by RT-qPCR (**Fig. 3A & 3B**). We also confirmed that, unlike the GAPDH mRNA, the endogenous GADD45B mRNA was not depleted upon transfection of KSHV SOX or HSV-1 vhs into 293T cells (**Fig. 3C**). Finally, we noted that siRNA-based depletion of GADD45B from iSLK.219 cells resulted in decreased efficiency of lytic KSHV reactivation as well as reduced expression of representative delayed early (ORF59) and late (K8.1) viral genes, suggesting that GADD45B expression is important for the KSHV lytic cycle (**Fig. S2**).

**Figure 3:**
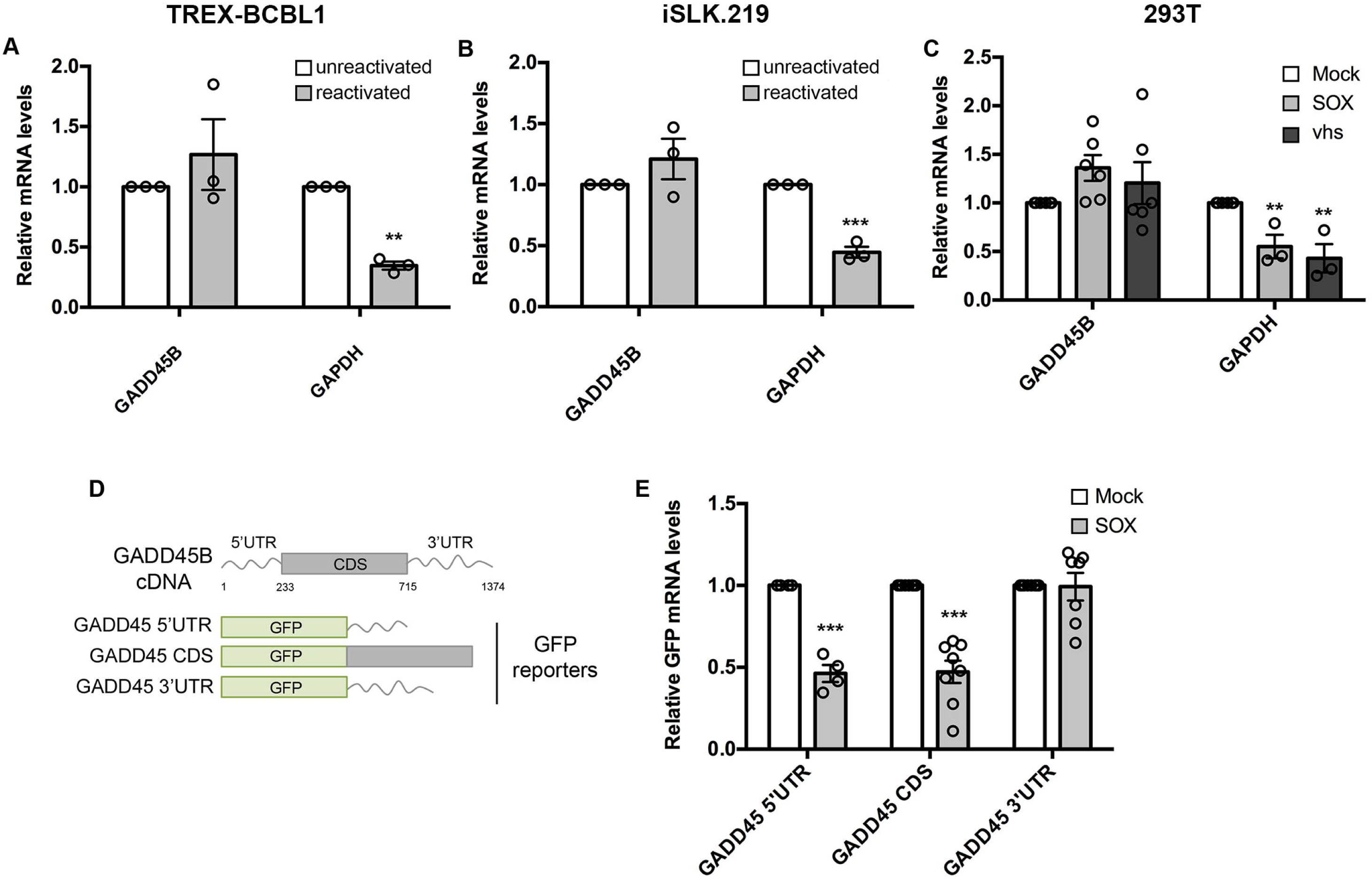
GADD45B mRNA is protected against SOX degradation. **(A, B)** Total RNA was extracted from unreactivated or reactivated KSHV-positive TREX-BCBL-1 (A) cells or iSLK.219 cells (B) and subjected to RT-qPCR to measure endogenous levels of the GADD45B or GAPDH transcript. **(C)** 293T cells were transfected with an empty vector or a plasmid expressing SOX or vhs. After 24 h, total RNA was harvested and subjected to RT-qPCR to measure endogenous levels of GADD45B or GAPDH transcripts. **(D)** Diagram of the reporter constructs where fragments of GADD45B transcript were fused to GFP. **(E)** 293T cells were transfected with the indicated GFP-GADD45B fusion constructs in the presence or absence of a plasmid expressing SOX. After 24 h, total RNA was harvested and subjected to RT-qPCR to measure GFP levels.

Recently, we showed that KSHV SOX cleaves its targets at a specific but degenerate RNA motif [21]. Thus, the failure of SOX (or perhaps vhs) to degrade the GADD45B mRNA could either be due to the absence of such a targeting motif (e.g. passive escape), or to the presence of a specific protective element like the IL-6 SRE (e.g. dominant escape). To distinguish these possibilities, we constructed chimeras between GFP, which has a well-characterized SOX cleavage element, and the GADD45B 5’ UTR, 3’ UTR, or coding region (CDS), each of which were cloned downstream of the GFP coding region (**Fig. 3D**). Co-transfection of the GFP-fused GADD45B 3’ UTR construct with SOX into 293T cells did not lead to degradation of this mRNA, whereas the GFP-GADD45B 5’ UTR or CDS fusions were readily degraded in SOX-expressing cells (**Fig. 3E**). Thus, similar to the IL-6 3’ UTR, the GADD45B 3’ UTR contains a protective sequence that prevents SOX cleavage of an established target mRNA.

### Hairpin structures within the GADD45B and IL-6 SREs are important for conferring resistance to SOX cleavage

To refine which GADD45B sequence encompassed the SRE, we initially looked for similarities between the GADD45B 3’ UTR and the IL-6 SRE using Clustal W alignment. While there were no stretches of significant sequence identity between the two RNAs, the last ∼200nt of the GADD45B 3’UTR had the highest similarity (∼46%) to the IL-6 SRE (**Fig. S3**). We therefore fused this putative SRE segment of the GADD45B 3’ UTR to GFP (GADD45B-SRE), and found that it was sufficient to confer nearly the same level of protection from SOX as the full GADD45B 3’ UTR in transfected 293T cells (**Fig. 4A**). Thus, similar to IL-6, the GADD45B 3’ UTR contains a ∼200 nt SRE (henceforth termed G-SRE).

**Figure 4:**
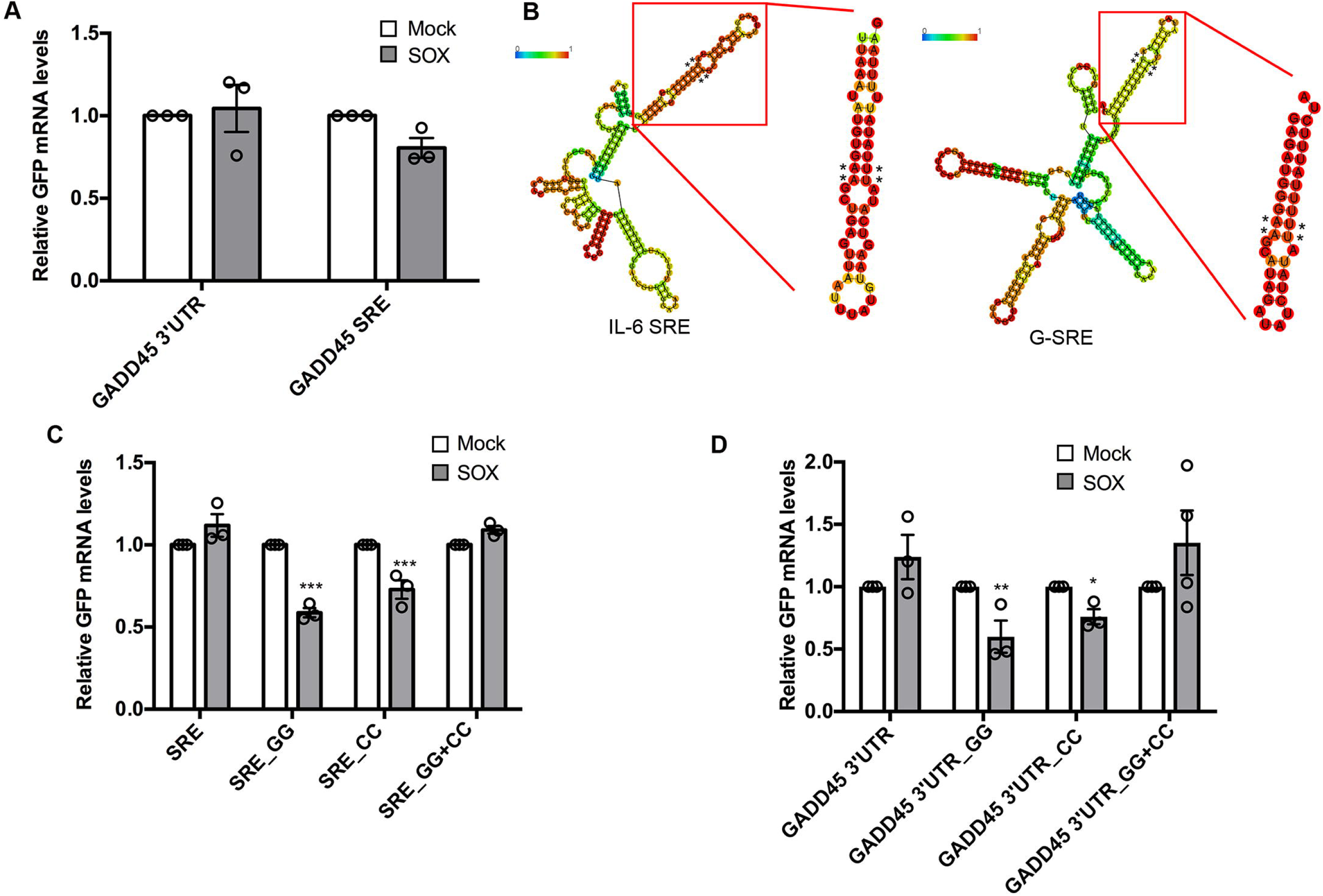
The SREs contain a long hairpin required for protection against SOX. **(A)** 293T cells were transfected with the indicated GFP reporter along with a control empty vector (mock) or a plasmid expressing SOX. After 24 h, total RNA was harvested and subjected to RT-qPCR to measure GFP mRNA levels. **(B)** Diagram of the structure prediction obtained with RNAfold for IL-6 and GADD45B SREs. The color scale represents the confidence score of the structure as calculated by RNAfold, with red representing the highest confidence. Asterisks denote the location of mutations that were introduced in the structure for the following assays. The insets are RNAfold predictions of the stem loops of interest in isolation. **(C)** 293T cells were transfected with the indicated GFP reporters containing mutations within IL-6 SRE at the residues marked by a * in (B) along with a control empty vector (mock) or a plasmid expressing SOX. After 24 h, total RNA was harvested and subjected RT-qPCR to measure GFP mRNA levels. **(D)** 293T cells were transfected with the indicated GFP reporters mutated within G-SRE at the residues marked by a * in (B) along with a control empty vector (mock) or a plasmid expressing SOX. After 24 h, total RNA was harvested and subjected to RT-qPCR to measure GFP mRNA levels.

We next sought to determine whether these two SREs could adopt a common secondary structure. RNAfold-based predictions showed that the 3’-most segment of both SREs form a long stem-loop protruding structure with at least one bulge near the middle of the stem, whereas the other regions of the SREs did not fold into any similar high-confidence structures (**Fig. 4B**). To validate this predicted SRE stem loop structure experimentally, we applied in-line probing, an RNA cleavage assay in which base-paired or structurally constrained nucleotides are protected from spontaneous phosphodiester bond hydrolysis [57]. Results from the cleavage reaction (**Fig. S4**) largely confirmed the RNAfold predictions, apart from a small variation in the IL-6 hairpin. We then tested whether this structure is required for either IL-6 SRE or G-SRE function by changing two conserved TT nucleotides located directly adjacent to the bulge in each hairpin structure to GG (SRE_GG; mutated residues marked with asterisks Fig. 4B). We also separately mutated the AA residues predicted to base pair with these nucleotides on the other side of the loop to CC (SRE_CC). Both the SRE_GG and the SRE_CC mutations in the IL-6 or GADD45B GFP fusions resulted in partial degradation of these mRNAs upon SOX expression in 293T cells, suggesting that disruption of that portion of the SRE hairpin impaired the protective capacity of each SRE **(Fig. 4C & 4D).** Notably, combining these mutations together (SRE_GG+CC), which should restore the secondary structure of the stem-loop, rescued the fully protective phenotype in SOX expressing cells (**Fig. 4C & 4D**). This suggests that, at least for this region of each SRE, RNA structure rather than the specific sequence is important for protection against SOX cleavage.

### The GADD45B and IL-6 SREs assemble a partially overlapping ribonucleoprotein complex

Using an *in vitro* RNA-pulldown based strategy, we previously determined that the IL-6 SRE assembles a specific ribonucleoprotein (RNP) ‘escape’ complex that is critical for its ability to mediate protection from cleavage by SOX [26,27]. Given the partial similarities in length, sequence, and structure between the IL-6 SRE and the G-SRE, we hypothesized that they might also assemble a similar set of RNPs to mediate their protective function. Indeed, it had already been established that both transcripts bind NCL [27,49]. We therefore performed Comprehensive Identification of RNA binding Proteins by Mass Spectrometry (ChIRP-MS) to compare the set of proteins bound *in vivo* to the IL-6 and GADD45B 3’UTRs in transfected 293T cells (**Fig. 5A**). Briefly, ChIRP-MS involves purifying an RNA of interest along with its associated proteins from crosslinked, sonicated cells using specific RNA probe-based capture, then identifying the bound proteins by MS [58]. Control probes directed against GFP were included to identify nonspecific interactions, which were then filtered out of the dataset. Using ChIRP-MS, we identified 195 proteins associated with the GADD45B 3’UTR and 245 proteins associated with the IL-6 3’UTR, of which 124 were in common between the two sets (**Table S1 & Fig. S5**). Many of the proteins previously identified as bound to the IL-6 SRE (**Fig. 5B**, gray nodes; [27]) were also recovered using ChIRP-MS for the IL-6 3’ UTR and were similarly associated with the GADD45B 3’ UTR (**Table S2** & **Fig. 5B**, purple nodes). We sought to independently validate a subset of these common interactors by western blot following ChIRP (**Fig. 5C**). Again, we recovered NCL and HuR (also known as ELAVL1), two proteins that were previously identified as important for IL-6 escape from SOX degradation [26,27]. We also confirmed hnRNPU, a protein that binds the IL-6 SRE but is not required for escape of the IL-6 SRE from SOX [27]. However, other IL-6 SRE-bound proteins required for its escape function were not detected on the GADD45B 3’ UTR by ChIRP-western blot, including IGF2BP1, STAU1, ZC3HAV1, YTHDC2, NPM1, and hnRNPD (**Fig. 5C)**. The fact that IGF2BP1 and NPM1 were recovered in the GADD45B and IL-6 ChIRP-MS experiments but not in the ChIRP-western blots likely reflects differences in sensitivity between MS and western blotting.

**Figure 5:**
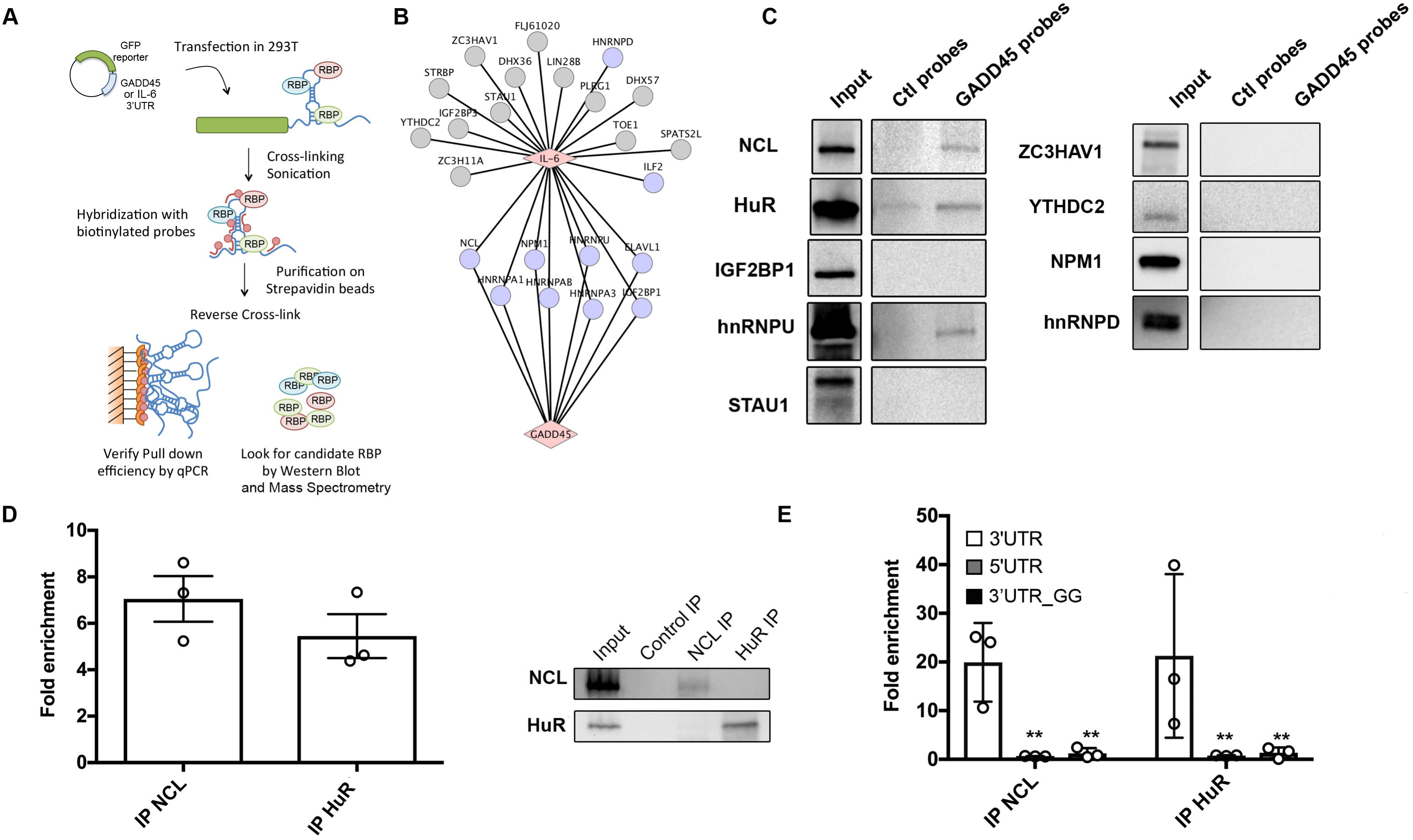
The IL-6 and GADD45B SRE bind a partially overlapping set of cellular proteins. **(A)** Diagram outlining the ChIRP assay. **(B)** Cytoscape network showing the proteins reported to interact with the IL-6 SRE from a previous screen [27] (gray nodes) overlaid with the set of proteins that were also recovered by ChIRP-MS for the IL-6 and GADD45B 3’ UTRs (purple nodes) **(C)** 293T were transfected with the GFP-GADD45B 3’UTR reporter, then 24h later they were subjected to ChIRP analysis and protein samples were western blotted. **(D)** Crosslinked lysates of 293T cells were subjected to RNA immunoprecipitation (RIP) with control IgG, anti-NCL, or anti-HuR antibodies and the level of co-purifying endogenous GADD45B mRNA was quantified by RT-qPCR. Bars represent the fold enrichment over the mock IP. **(E)** 293T cells were transfected with either the GFP-GADD45B 5’UTR, GFP-GADD45B 3’UTR, or GFP-GADD45B 3’UTR_GG reporter for 24 h. Lysates were then subjected to RIP as described in (C). Bars represent the fold enrichment over mock IP.

To further validate the interactions with NCL, HuR, and hnRNPU, we immunoprecipited (IP) each endogenous protein from 293T cells and performed RT-qPCR to measure the level of co-precipitating endogenous GADD45B RNA. We observed a >5-fold enrichment of GADD45B mRNA over the mock (IgG) IP for both NCL and HuR, although we were not able to detect an association with hnRNPU in this assay (data not shown) (**Fig. 5D**). We confirmed that the interaction occurs on the GADD45B 3’UTR by performing the IPs from cells transfected with a GFP reporter fused to either the GADD45B 5’ UTR or 3’ UTR (**Fig. 5E**). The structurally compromised mutant GADD45B 3’UTR_GG described in Fig. 4B failed to interact with NCL and HuR, confirming that the stem-loop structure in the SRE is important for protein binding (**Fig. 5E**). Collectively, these data suggest that while there is some overlap between the sequence, structure, and RNA binding proteins associated with the GADD45B and IL-6 SREs, these elements likely assemble distinct RNP complexes and thus may function in a related but non-identical manner.

### The GADD45B and IL-6 SREs share a requirement for HuR but differ in the breadth of their protective capacity

Depletion of either HuR or NCL impairs the ability of the IL-6 derived SRE to protect its associated mRNA from degradation by SOX [26,27]. Given that both proteins are also bound by the G-SRE, we evaluated whether they were similarly important for G-SRE-mediated escape. We individually depleted each protein from 293T cells using siRNAs targeting HuR or NCL (or control non-targeting siRNAs), then measured the ability of SOX to degrade the GFP-GADD45B-3’UTR reporter by RT-qPCR (**Fig. 6A & 6B**). Similar to the IL-6 SRE, depletion of HuR eliminated the protective capacity of the G-SRE, leading to degradation of the GFP-GADD45B-3’UTR mRNA in SOX-expressing cells (**Fig. 6A**). Surprisingly however, depletion of NCL did not impair the protective effect of the G-SRE in SOX-expressing cells (**Fig. 6B**).

**Figure 6:**
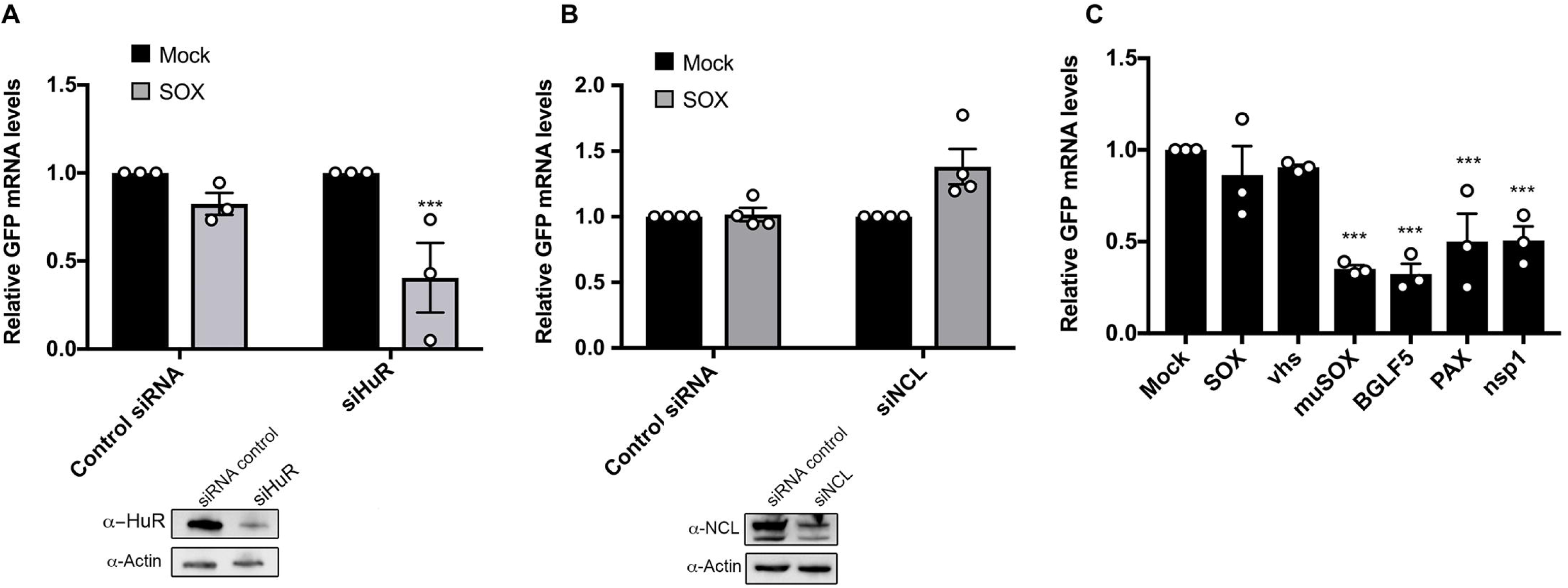
The GADD45B SRE function requires HuR, but is only protective against SOX and vhs. (**A, B**) 293T cells were treated with siRNAs targeting HuR (A) or NCL (B) or control non targeting siRNAs. 48h later, cells were transfected with the GFP-GADD45B-3’UTR reporter along with a control empty vector or a plasmid expressing SOX. After 24 h, total RNA was harvested and subjected to RT-qPCR to measure GFP mRNA levels. Protein levels of HuR and NCL after siRNA-mediated depletion are shown under the bar graphs (∼30% and 45% of total protein left respectively) **(C)** 293T cells were transfected with the GFP-GADD45B-3’UTR reporter along with a control empty vector or a plasmid expressing the indicated viral endonucleases. After 24 h, total RNA was harvested and subjected to RT-qPCR to measure GFP mRNA levels.

Finally, we used the GFP-GADD45B-3’UTR reporter to evaluate whether the G-SRE conferred protection from cleavage by the panel of viral endonucleases described in Fig. 1. Using the same experimental set up as was used for the IL-6 derived SRE reporter, we observed that while the GADD45B 3’ UTR protected against SOX and vhs, it was unable to protect against degradation by muSOX, BGLF5, and PA-X (**Fig. 6C**). Furthermore, it did not protect against cleavage by the nsp1-activated host endonuclease (**Fig. 6C**). Thus, although the IL-6 and GADD45B derived nuclease escape elements exhibit a number of similarities, they are distinct in both their RNP complex requirements and in the breadth of nucleases they restrict.

## DISCUSSION

Viruses extensively interface with the host gene expression machinery to promote their own RNA and protein synthesis and to control the cellular response to infection. In this regard, they have proven to be invaluable tools to dissect mechanisms of gene regulation. Here, we reveal that a ∼200 nt sequence present in the 3’ UTR of the cellular IL-6 mRNA functions as a broad-acting, virus-specific endonuclease escape element, and identify a similar element in the GADD45B 3’ UTR. Although these two SREs are not identical in their protective capacity, they display some sequence and putative structural similarities and are both functionally dependent on HuR binding. We hypothesize that these may therefore be representative members of a new type of RNA regulatory element engaged during viral endonuclease-triggered mRNA decay. Determining whether other such elements exist in mammalian or viral mRNAs is an important future goal, as is deciphering conditions under which such elements impact RNA fate in uninfected cells.

The IL-6 and GADD45B SREs display relatively limited sequence similarity but have at least one structurally important stem-loop in common. Thus, RNA structure appears to be a central component of an SRE and is likely to influence recruitment or arrangement of proteins involved in escape. This is in line with the fact that RNP assembly can be more heavily impacted by structural fidelity than primary sequence recognition [59]. We hypothesize that there is a core set of proteins required for SRE function, such as HuR, but that individual SREs recruit distinct accessory factors that dictate the conditions or mechanism by which that SRE protects against nuclease targeting. In this regard, we have not found significant overlap between known HuR mRNA targets and mRNAs that are not downregulated by SOX [22]. This is in agreement with the fact that many of the SRE bound proteins have a breadth of roles in controlling RNA fate beyond conferring nuclease escape [60-64]. SRE activity may be further impacted by the cellular context and neighboring regulatory features, particularly in the case of tightly controlled transcripts. The 3’ UTR of IL-6, for example, is targeted by several cellular endonucleases including MCPIP1 [65-67], which has recently been reported to be inhibited during de novo KSHV infection [68]. The above features underscore the importance of characterizing multiple SREs to parse out specific protein requirements, but also present challenges for the identification of SREs, as they cannot be easily predicted. Indeed, it has taken more than a decade from the first report of IL-6 escaping SOX-induced host shutoff to identify a second SRE-containing mRNA [29].

The observation that the IL-6 SRE in particular acts against diverse viral endonucleases but does not impact host endonucleases has important implications for understanding target recognition by these host shutoff factors. First, it suggests that there are key commonalities to how the viral endonucleases are recruited to mRNAs, and that these features are distinct from those involved in cellular endonuclease recruitment. The viral endonucleases recognize translation competent mRNAs, but do not require ongoing translation for cleavage [10]. While vhs is recruited to a 5’ cap proximal location due to its interaction with a eIF4H, the SOX homologs do not respond to particular location cues on their target mRNA, and instead recognize a degenerate target motif that can be positioned anywhere [10,21,69-71]. PA-X has been suggested to bind RNA processing factors in the nucleus, but how it recognizes cytoplasmic mRNA targets remains unknown [72]. By contrast, the cellular NMD and no-go decay quality control pathways show a clear dependence on translation for target recognition, and mRNA cleavage occurs at the site of the error or translation stall [13]. The observation that neither the IL-6 nor GADD45B SREs impede these cellular endonucleases indicates that SREs are not inhibitory to translation. Furthermore, our FISH data do not support mRNA relocalization or sequestration as a driving feature of SRE-mediated escape. We instead hypothesize that the SRE somehow occludes one or more factors required for recruitment of the viral endonucleases. This could occur via long-range interactions with mRNA cap-associated proteins, which we previously showed can take place for the IL-6 SRE [27]. The fact that the GADD45B SRE restricts against a more limited set of host shutoff factors argues against a single factor requirement. However, it is possible that there are multiple binding sites on one factor that are more comprehensively blocked by the IL-6 SRE. Independent of the mechanisms involved, nuclease escape elements could be developed as tools to broadly inhibit viral endonucleases without disrupting normal RNA decay pathways.

How might viruses benefit from maintaining the expression of IL-6 and GADD45B during infection? The requirement for IL-6 during KSHV infection is well documented [30-38], but the role of GADD45B may be more nuanced. While we demonstrated that it escapes host shutoff during lytic KSHV infection and appears important for viral reactivation in the iSLK.219 model, GADD45B expression is repressed by viral miRNAs during latency to avoid its cell cycle arrest and pro-apoptotic functions [73]. However, GADD45B has a number of additional activities that could be necessary for the viral lytic cycle. These include promoting DNA demethylation and Retinoblastoma (Rb) inactivation, processes that are important for a number of DNA viruses, including KSHV [74-79]. If our hypothesis that there are numerous SREs throughout the transcriptome is correct, it is likely that individual viruses require only a subset of these host genes for replication. Other mRNAs may collaterally escape simply because they contain SREs that functionally mimic those present in the ‘required’ mRNAs. In this regard, the fact that a given mRNA escapes degradation by multiple viral endonucleases does not necessarily indicate that expression of that gene is broadly required for infection. Finally, it is important to bear in mind that the SREs discovered thus far are cellular elements that assemble cellular proteins—and thus presumably play host-directed roles in the regulation of their associated transcripts. It may therefore be the case that some SRE-bearing mRNAs function in an antiviral capacity. Exploring possible virus-host evolutionary interplay for SREs remains an exciting prospect for future studies.

## MATERIALS AND METHODS

### Cells and transfections

The KSHV-positive B cell line bearing a doxycycline-inducible version of the major lytic transactivator RTA (TREX-BCBL-1) [55] was maintained in RPMI medium (Invitrogen) supplemented with 10% fetal bovine serum (FBS; Invitrogen), 200 μM L-glutamine (Invitrogen), 100 U/ml penicillin/streptomycin (Invitrogen), and 50 μg/ml hygromycin B (Omega Scientific). Lytic reactivation was induced by treatment with 20 ng/ml 2-O-tetradecanoylphorbol-13-acetate (TPA; Sigma), 1 μg/ml doxycycline (BD Biosciences), and 500 ng/ml ionomycin (Fisher Scientific) for 48h. 293T cells (ATCC) were grown in DMEM (Invitrogen) supplemented with 10% FBS. The KHSV-infected renal carcinoma cell line iSLK.219 bearing doxycycline-inducible RTA was grown in DMEM supplemented with 10% FBS [56]. KSHV lytic reactivation of the iSLK.219 cells was induced by the addition of 0.2 μg/ml doxycycline (BD Biosciences) and 110 μg/ml sodium butyrate for 48 h.

For DNA transfections, cells were plated and transfected after 24h when 70% confluent using linear PEI (polyethylenimine). For small interfering RNA (siRNA) transfections, 293T cells were reverse transfected in 12-well plates by INTERFERin (Polyplus-Transfection) with 10 μM of siRNAs. siRNAs were obtained from IDT as DsiRNA (siRNA SMG6: hs.Ri.SMG6.13.1; siRNA NCL: hs.Ri.NCL.13.1; siRNA HuR ELAVL1#1: hs.Ri.ELAVL1.13.2; siRNA ELAVL1#2: hs.Ri.ELAVL1.13.3). 48h following siRNA transfection, the cells subjected to DNA transfection as indicated.

### Plasmids

The GADD45B 5’UTR and CDS were obtained as G-blocks from IDT and cloned into a pcDNA3.1 plasmid downstream of the GFP coding sequence. GADD45B 3’UTR was cloned from a pDest-765 plasmid (kindly provided by J. Ziegelbauer) into a pcDNA3.1 plasmid downstream of the GFP coding sequence. The GFP-IL-6 3’UTR, SRE and ΔSRE fusion constructs were described previously [27]. The dsRed2 and dsRed2-PTC reporters were described elsewhere [10]. The SRE and ΔSRE were PCR amplified from the GFP reporters and cloned downstream of the dsRed2 ORF.

Point mutations were introduced with the Quickchange site directed mutagenesis protocol (Agilent) using the primers described in **Table S3**.

### FiSH

Stellaris FiSH probes recognizing MS2 and GAPDH labeled with Quasar 570 Dye (MS2: SMF-1063-5; GAPDH: SMF-2026-1) were hybridized in 293T cells following manufacturer’s instructions available online at www.biosearchtech.com/stellarisprotocols. Briefly, 293T cells were grown on coverslips and transfected with either an empty vector control or the MS2-SRE or MS2ΔSRE constructs. 24h later, cells were washed, fixed in 4% formaldehyde and permeabilized in 70% ethanol. Probes (12.5µM) were then hybridized for >5h at 37°C in Vanadyl ribonucleoside (10mM), formamide (10%), saline sodium citrate (SSC), dextran (10%) and BSA (0.2%). DAPI was added for the last hour to stain cell nuclei. Coverslips were washed in SSC and mounted in Vectashield mounting medium (VectorLabs) before visualization by confocal microscopy on a Zeiss LSM 710 AxioObserver microscope.

### Comprehensive Analysis of RNA Binding Proteins (ChIRP)

ChIRP was carried out according to a protocol published previously [58] with minor modifications. Briefly, 293T cells were transfected with the GFP-GADD45B 3’UTR plasmid and 24h later cells were crosslinked in 3% formaldehyde for 30 minutes, quenched in 125mM Glycine, washed with PBS and the pellet was flash frozen. Pellets were then lysed in fresh ChIRP Lysis Buffer [Tris pH 7.4 50mM, EDTA 10mM, SDS 1%], sonicated, and cell debris was removed by centrifugation at (14,000rpm for 10 min). Two mL of hybridization buffer was added to each sample along with 100pmol of the indicated probes (see **Table S3** for sequences). Hybridization was carried out for 4-12h. After adding streptavidin beads to the samples for 1h, samples were washed and reverse crosslinked at 65°C overnight. For western blotting, beads were resuspended in 4X loading buffer, boiled for 30 minutes and resolved by SDS-PAGE. For mass spectrometry, recovered proteins from two independent biological replicates were gel extracted and subjected to LC-MS/MS at the UC Berkeley Vincent J. Coates Proteomics/Mass Spectrometry Laboratory.

### In-line probing

RNA was synthesized by Dharmacon (SRE-HP #CTM-299208, GADD45- HP- #CTM-299209). Approximately 1ug of RNA was 5’ end-labeled and purified on an 8M urea gel. Samples were then ethanol precipitated in presence of glycogen, washed in 70% ethanol and dissolved in 20 µL of DEPC-treated water. Reactions were carried out with ∼1uL of purified RNA (∼100,000cpm). In-line probing was performed following a standard procedure [57]. Briefly, in-line probing assays were carried for 40h in 2X in-line buffer [100 mM Tris·HCl, pH 8.3, 40 mM MgCl2, 200 mM KCl] and quenched by adding the same volume of RNA loading Buffer [95% formamide, 10 mM EDTA and 0.025% xylene cyanol]. The no-reaction (NR) treatment, RNase T1 (T1) and partial base hydrolysis (–OH) ladders were prepared as 20µL reactions and quenched with 20 µL RNA loading buffer. Dried gels were exposed on a phosphorimager screen and scanned using a Typhoon laser-scanning system (GE Healthcare).

### RT-qPCR

Total RNA was harvested using Trizol following the manufacture’s protocol. cDNAs were synthesized from 1 µg of total RNA using AMV reverse transcriptase (Promega), and used directly for quantitative PCR (qPCR) analysis with the DyNAmo ColorFlash SYBR green qPCR kit (Thermo Scientific). Signals obtained by qPCR were normalized to 18S.

### RNA Immunoprecipitation

Cells were crosslinked in 1% formaldehyde for 10 minutes, quenched in 125mM glycine and washed in PBS. Cells were then lysed in low-salt lysis buffer [NaCl 150mM, NP-40 0.5%, Tris pH8 50mM, DTT 1mM, MgCl2 3mM containing protease inhibitor cocktail and RNase inhibitor] and sonicated. After removal of cell debris, specific antibodies were added as indicated overnight at 4°C. Magnetic G-coupled beads were added for 1h, washed three times with lysis buffer and twice with high-salt lysis buffer (low-salt lysis buffer except containing 400mM NaCl). Samples were separated into two fractions. Beads containing the fraction used for western blotting were resuspended in 30μL lysis buffer. Beads containing the fraction used for RNA extraction were resuspended in Proteinase K buffer (NaCl 100mM, Tris pH 7.4 10mM, EDTA 1mM, SDS 0.5%) containing 1μL of PK (Proteinase K). Samples were incubated overnight at 65°C to reverse crosslinking. Samples to be analyzed by western blot were then supplemented with 10μL of 4X loading buffer before resolution by SDS-PAGE. RNA samples were resuspend in Trizol and were processed as described above.

### Western Blotting

Lysates were resolved by SDS-PAGE and western blotted with the following antibodies at 1:1000 in TBST (Tris-buffered saline, 0.1% Tween 20): rabbit anti- EST1A/SMG6 (Abcam), rabbit anti-NCL (Abcam), rabbit anti-HuR (Millipore), rabbit anti-actin (Abcam), rabbit anti-IGF2BP1 (Abcam), rabbit anti-YTHDC2 (Abcam), rabbit anti-hnRNPU (Abcam), rabbit anti-ZC3HAV1 (Abcam), rabbit anti-hnRNPD (Abcam), rabbit anti- STAU1 (Pierce), mouse anti-NPM1 (Abcam), rabbit anti-GADD45 (Abcam), mouse anti-GAPDH (Abcam), mouse anti-ORF59 (Adv biotechnologies), or rabbit anti-K8.1 (PRF&L, inc). Primary antibody incubations were followed by HRP-conjugated goat anti-mouse or goat anti-rabbit secondary antibodies (Southern Biotechnology, 1:5000).

### Statistical analysis

All results are expressed as means ± S.E.M. of experiments independently repeated at least three times. Unpaired Student’s t test was used to evaluate the statistical difference between samples. Significance was evaluated with P values as follows: * p<0.1; ** p<0.05; *** p<0.01.

## ACKNOWLEDGMENTS

We thank members of the Glaunsinger and Coscoy labs for helpful discussions. We also thank Ming Hammond, Debojit Bose and Carolin Vogt for technical advice on in-line probing.

## FUNDING

This research was supported by NIH grants CA160556 and CA136367 (http://www.nih.gov/) and a Burroughs Wellcome Investigator in the Pathogenesis of Infectious Diseases Award (http://www.bwfund.org/grant-programs/infectious-diseases/investigators-in-pathogenesis-of-infectious-disease) to BAG. B.A.G. is an investigator of the Howard Hughes Medical Institute. Funding for open access charge: Howard Hughes Medical Institute and MM is a HHMI Fellow of the Damon Runyon Cancer Research Foundation (DRG- 2207-14) (http://www.damonrunyon.org/). This work used the Vincent J. Proteomics/Mass Spectrometry Laboratory at UC Berkeley, supported in part by NIH S10 Instrumentation Grant S10RR025622.

## SUPPORTING INFORMATION

**Figure S1:** 293T cells were transfected with the indicated MS2 luciferase reporter containing or lacking the IL-6 SRE +/- a KSHV SOX expression plasmid. After 24 h, total RNA was harvested and subjected RT-qPCR to measure reporter mRNA levels.

**Figure S2: (A)** iSLK.219 cells were treated with siRNAs targeting GADD45B (or control non-target siRNAs) for 48h. Cells were then reactivated with doxycycline and sodium butyrate for 24 or 48h, fixed, and reactivation efficiency was monitored by expression of red fluorescent protein, which is expressed from the viral genome under the control of the lytic PAN promoter. **(B)** After siRNA treatment and reactivation as described in A, cell lysates were subjected to western blotting to measure protein levels of GADD45B, the KSHV proteins K8.1 and ORF59, and GAPDH (as a loading control).

**Figure S3:** Clustal W alignment of IL-6 SRE and GADD45B 3’UTR.

**Figure S4:** In-line probing of the IL-6 (left) and GADD45B (right) predicted stem-loop. ^32^P- labeled RNA (NR, no reaction) and products resulting from partial digestion with nuclease T1 (T1; cuts after G residues), partial digestion with alkali (-OH), and spontaneous cleavage during a 24h incubation are shown. Product bands corresponding to G residues (generated by T1 digestion) are labeled with black arrows. Predicted paired or unpaired residues are marked on the right of each gel and are shown on the RNA fold diagrams.

**Figure S5:** Network representation of the full set of proteins identified by ChIRP-MS for either the IL-6 or GADD45B 3’UTR. Purple nodes represent proteins that were previously identified using an *in vitro* pulldown/MS-based assay [27].

**Table S1:** List of proteins identified by ChIRP-MS. Proteins in bold were in common between the IL-6 and GADD45B 3’ UTR datasets. Background proteins found in the control ChIRP-MS runs with probes were filtered out together with common contaminants.

**Table S2**: List of proteins identified by ChIRP-MS that are in common with previously identified SRE-binding proteins [27].

**Table S3:** List of primers used in this study

